# Expression of ACE2 and TMPRSS2 proteins in the upper and lower aerodigestive tracts of rats

**DOI:** 10.1101/2020.05.14.097204

**Authors:** Taku Sato, Rumi Ueha, Takao Goto, Akihito Yamauchi, Kenji Kondo, Tatsuya Yamasoba

**Affiliations:** Department of Otolaryngology and Head and Neck Surgery, Faculty of Medicine, the University of Tokyo, Tokyo, Japan

**Keywords:** ACE2, TMPRSS2, COVID-19, dysgeusia, peripheral lung

## Abstract

**Objective:** Patients with coronavirus disease 2019 (COVID-19), caused by severe acute respiratory syndrome coronavirus 2 (SARS-CoV-2), exhibit not only respiratory symptoms but also symptoms of chemo-sensitive disorders and kidney failure. Cellular entry of SARS-CoV-2 depends on the binding of its spike protein to a cellular receptor named angiotensin-converting enzyme 2 (ACE2), and the subsequent spike protein-priming by host cell proteases, including transmembrane protease serine 2 (TMPRSS2). Thus, high expression of ACE2 and TMPRSS2 are considered to enhance the invading capacity of SARS-CoV-2.

**Methods:** To elucidate the underlying histological mechanisms of the aerodigestive disorders caused by SARS-CoV-2, we investigated the expression of ACE2 and TMPRSS2 proteins in the aerodigestive tracts of the tongue, hard palate with partial nasal tissue, larynx with hypopharynx, trachea, esophagus, lung, and kidney of rats through immunohistochemistry.

**Results:** Strong co-expression of ACE2 and TMPRSS2 proteins was observed in the nasal respiratory epithelium, trachea, bronchioles, alveoli, kidney, and taste buds of the tongue. Remarkably, TMPRSS2 expression was much stronger in the peripheral alveoli than in the central alveoli. These results coincide with the reported clinical symptoms of COVID-19, such as the loss of taste, loss of olfaction, respiratory dysfunction, and acute nephropathy.

**Conclusions:** A wide range of organs have been speculated to be affected by SARS-CoV-2 depending on the expression levels of ACE2 and TMPRSS2. Differential distribution of TMPRSS2 in the lung indicated the COVID-19 symptoms to possibly be exacerbated by TMPRSS2 expression. This study might provide potential clues for further investigation of the pathogenesis of COVID-19.

**Level of Evidence:** NA

## Introduction

The coronavirus disease 2019 (COVID-19), driven by the novel severe acute respiratory syndrome coronavirus 2 (SARS-CoV-2), is explosively spreading worldwide since its first report in December 2019 ^1^. Viral loads of SARS-CoV-2 have been found to be high in the upper respiratory tract, especially in the nasopharynx ^2^, whereas that of SARS-CoV had been reported to be high in the lower respiratory tract ^3^.

The symptoms of COVID-19 in patients are mostly similar to those of other respiratory infections, including that of SARS: fever (43–98%), cough (68–82%), fatigue (38– 44%), sore throat (14–17%), and sputum (28–33%) ^4,5^. Moreover, dysfunction of senses, such as loss or change of taste (ageusia or dysgeusia) and loss of smell (anosmia), has been consistently reported as a unique clinical feature of this disease (19–74%) ^6,7^.

Angiotensin-converting enzyme 2 (ACE2) is known to be the receptor responsible for the cellular entry of coronaviruses, including SARS-CoV-2 ^8,9^. Once the viral spike (S) protein binds to ACE2, it is primed by the transmembrane protease serine 2 (TMPRSS2) of host cells, thereby facilitating viral entry ^10^. Thus, high expression of ACE2 and TMPRSS2 may be considered to enhance the invasion of SARS-CoV-2. Previous genetic studies had demonstrated high expression of ACE2 on the human oral mucosa, including tongue epithelium ^11^, and co-expression of ACE2 and TMPRSS2 in both human nasal and bronchial epithelia ^12^. However, histological studies have been limited.

Based on the aforementioned aspects, this study aimed to elucidate the underlying histological mechanisms of the upper and lower aerodigestive disorders by studying the expression of ACE2 and TMPRSS2 on the aerodigestive tracts of rats.

## Materials and Methods

### Rat tissue samples

Tissue samples were obtained from rats examined in previous studies ^13,14^. Five eight-week-old male Sprague Dawley rats were included in the control group. The following paraffin-embedded tissues were collected: the tongue, hard palate with partial nasal tissue, larynx with hypopharynx, trachea, esophagus, lung, and kidney. All animal experiments were conducted in accordance with institutional guidelines and with the approval of the Animal Care and Use Committee of the University of Tokyo (No. P14-051, P17-126).

### Histological analyses

To detect the expression of ACE2 and TMPRSS2 in the upper and lower airway, histological analysis was performed by immunostaining. Four-micron-thick serial paraffin sections were deparaffinized in xylene and dehydrated in ethanol before immunostaining.

Prior to immunostaining, deparaffinized sections were treated with 3% hydrogen peroxide to block endogenous peroxidase activity, and then incubated in Blocking One (Nacalai Tesque, Kyoto, Japan) to block any non-specific binding to immunoglobulin. After antibody activation, primary antibodies against ACE2 (1:300 dilution; rabbit monoclonal, Abcam, ab108252; Cambridge, UK) and TMPRSS2 (1:1000 dilution; rabbit monoclonal, Abcam, ab92323; Cambridge, UK) were detected with appropriate peroxidase-conjugated secondary antibodies and a diaminobenzidine substrate. Images of all sections were captured using a digital microscope (Keyence BZ-X700) with 4× and 20× objective lenses.

## Results

In the tongue, both ACE2 and TMPRSS2 proteins were strongly co-expressed in the taste buds of foliate papillae (Fig.1A, B). In the nasal cavity, the olfactory neuroepithelium showed sparse ACE2-positive cells, and remarkable TMPRSS2 immunostaining in the cytoplasm. In the villous brush border of the respiratory columnar epithelium, ACE2 and TMPRSS2 were highly co-expressed (Fig.1C, D). In the palate, ACE2 and TMPRSS2 were weakly and diffusely co-expressed in the squamous epithelium (Fig. 1E).

**Figure 1.**
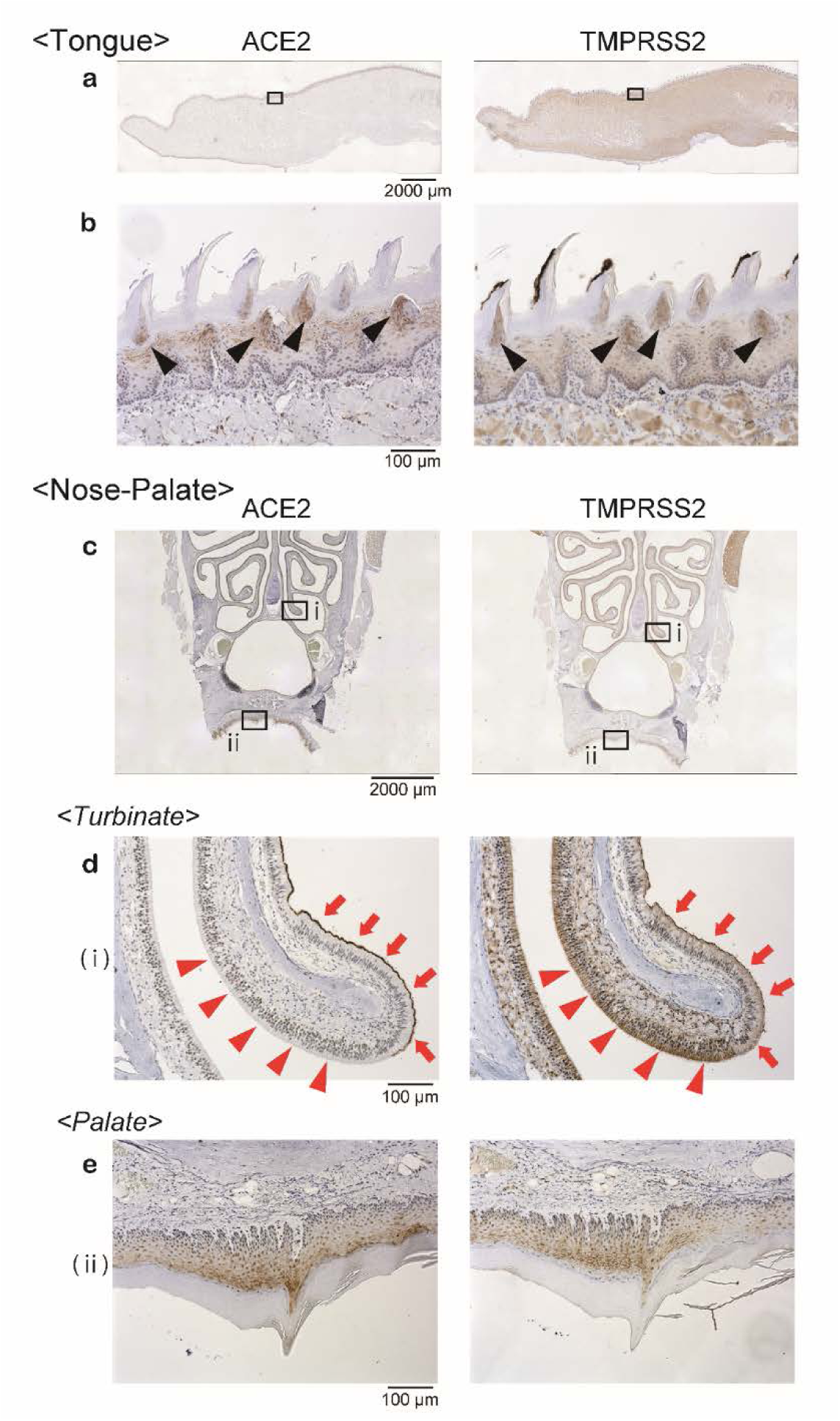
Representative images of tissues, stained with antibodies against ACE2 and TMPRSS2, are shown. a and b: Taste buds of foliate papillae in the tongue were strongly stained for ACE2 and TMPRSS2 (arrow heads) (a, magnification, 40×; b, magnification, 200×). c: An inferior part of the nasal tissue and palate are shown. The boxes indicate a part of the turbinate (i) and palate (ii), both shown at higher magnification (magnification, 200x) in (d) and (e). d: In the olfactory epithelium, ACE2-positive cells were sparse (red arrow heads) while TMPRSS2-positive cells were abundant with strong cytoplasmic expression (red arrow heads). In the nasal respiratory columnar epithelium, while ACE2 was strongly positive in the cilia (red arrows), TMPRSS2 was weakly positive in the cytoplasm (red arrows). e:Epithelial cells of the palate showed diffuse immunopositivity for ACE2 and TMPRSS2.

In the epithelium of the larynx, TMPRSS2 expression was explicit, whereas ACE2 staining was weak and sparse. This tendency was similarly observed in the whole larynx from epiglottis to the subglottis (Fig. 2). In the trachea, the cilia of primary tracheal epithelial cells showed high expression of ACE2 and TMPRSS2. In addition, TMPRSS2 was abundantly present in the cytoplasm of ciliated columnar epithelium (Fig. 3A, B). In the lung, consistent with findings in the trachea, ACE2 and TMPRSS2 were strongly expressed in the epithelium of bronchiole, in alveolar epithelial cells, and in capillary endothelium. Remarkably, TMPRSS2-positive cells were more abundant in the peripheral alveoli than in the central part of the lung (Fig. 4A, B).

**Figure 2.**
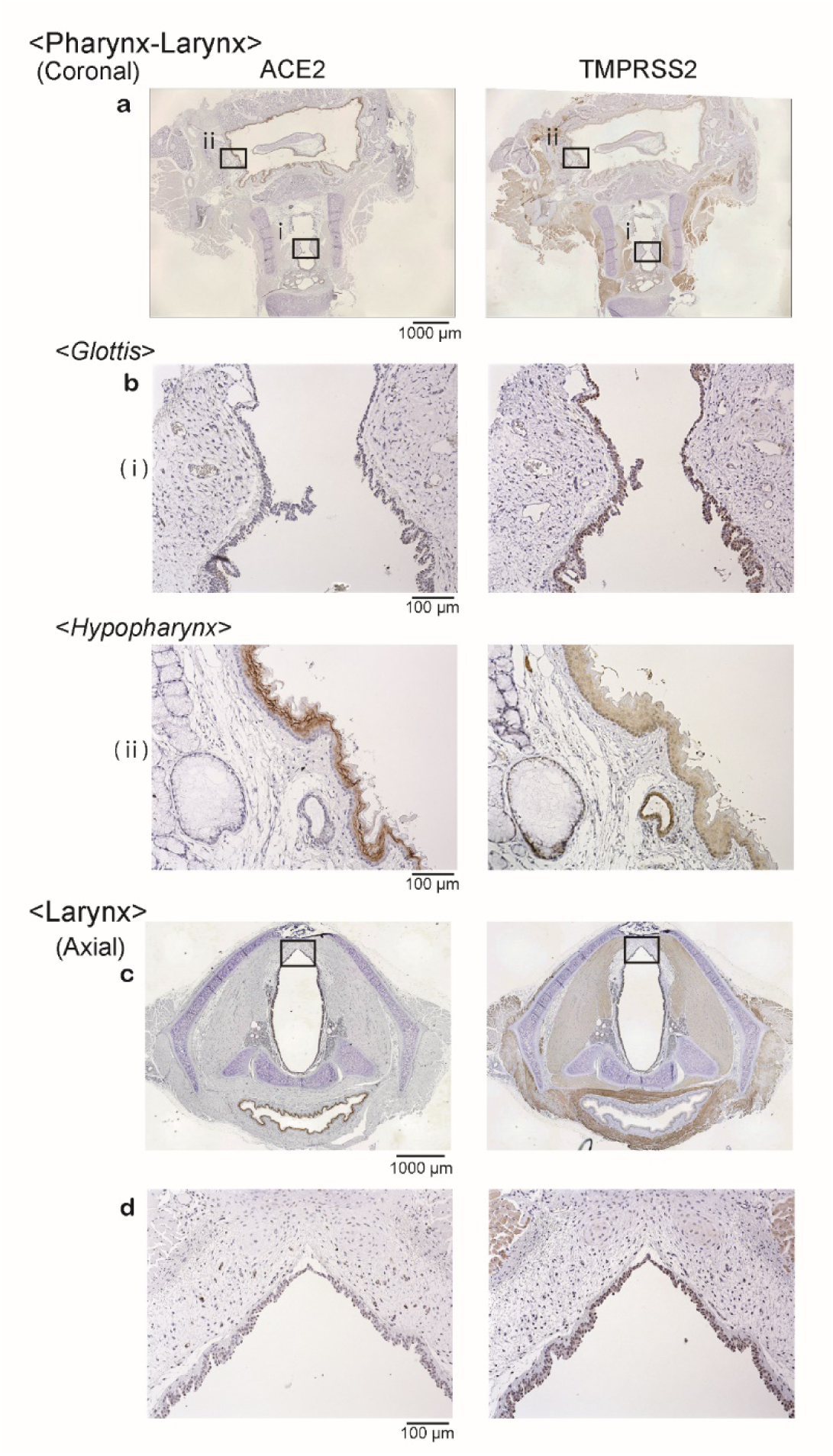
Representative images of the larynx and hypopharynx, stained with antibodies against ACE2 and TMPRSS2, are shown. **a**: Coronal images of the larynx and pharynx at lower magnification (magnification, 40×). The boxes indicate regions of the glottis (i) and hypopharynx (ii). **b**: In the epithelium of the glottis, ACE2-positive cells were observed slightly while TMPRSS2-positive cells were explicit. In the epithelium of hypopharynx, ACE2 was expressed strongly, whereas TMPRSS2 was expressed diffusely. **c**: Axial images of the larynx at lower magnification (magnification, 40×). **d**: The boxes in (c) indicate the regions of anterior macula flava, which are shown at higher magnification (magnification, 200x) in (d). ACE2-positive cells were detected only slightly in the epithelium and sub-epithelial tissue, and moderately TMPRSS2-positive cells were detected in the epithelium.

**Figure 3.**
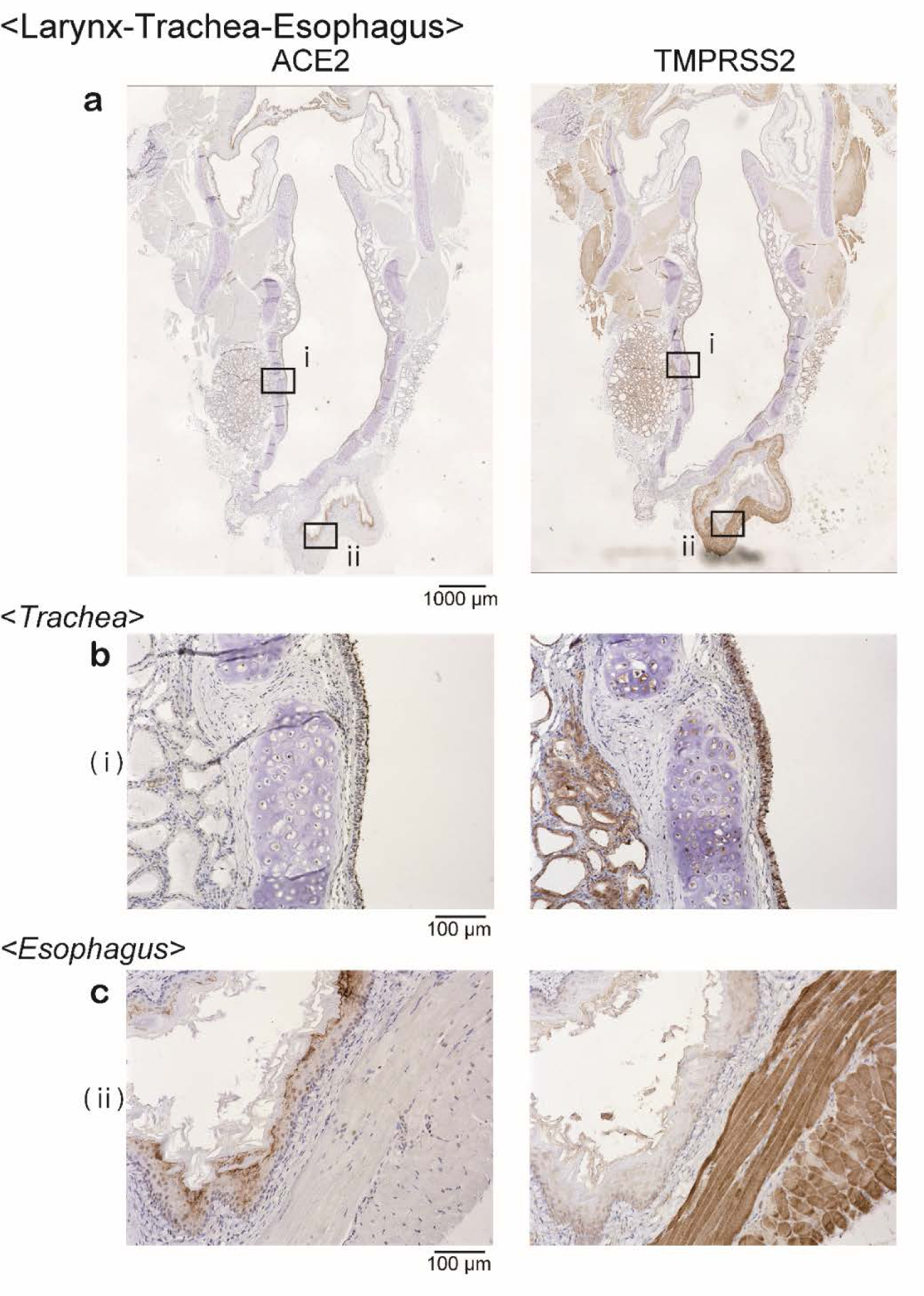
Representative images of the trachea and esophagus, stained with antibodies against ACE2 and TMPRSS2, are shown. **a**: Low-power field micrographs of coronal sections (magnification, 40×). Each box (i, ii) in (a) indicates the region of trachea and esophagus, respectively, shown at a representative higher magnification (magnification, 200x) in (b, c). **b**: In the trachea, cilia of the epithelial cells were weakly stained for ACE2 while the cells were strongly stained for TMPRSS2. **c**: In the esophagus, the surface of squamous epithelium was distinctly stained for ACE2. Positive staining for TMPRSS2 was weakly present in the cytoplasm of the epithelium and clearly so in the esophageal muscles.

**Figure 4.**
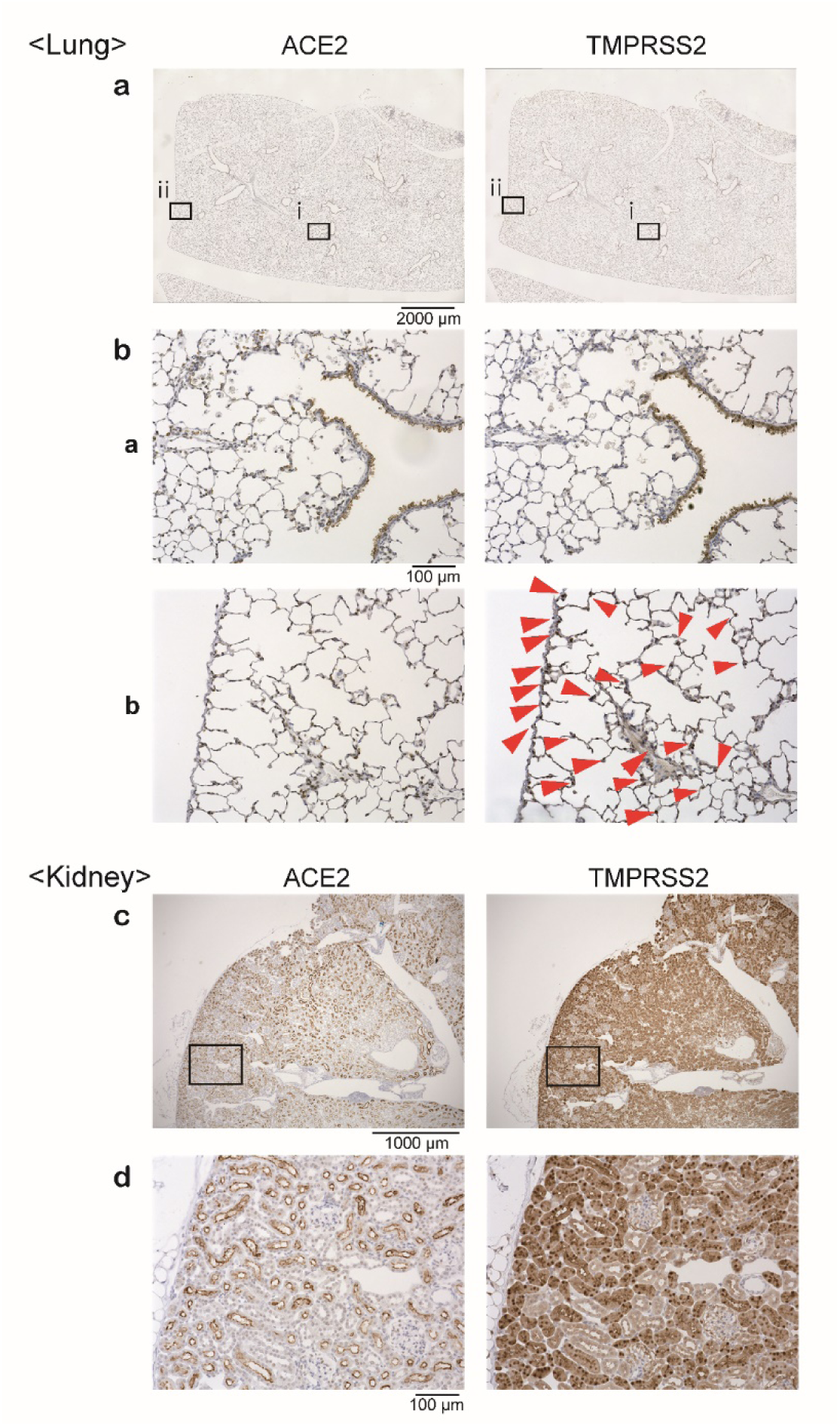
Representative images of the lung and kidney, stained with antibodies against ACE2 and TMPRSS2, are shown. a: Low-magnification images of the lung (magnification, 40×). The boxes (i, ii) indicate the central (i) and peripheral parts (ii) of the lung, respectively. b: High-magnification images of the lung (magnification, 200x). TMPRSS2 was more strongly expressed in the peripheral alveoli (red arrow heads) compared to that in the central part of the lung. c, d: Representative images of the kidney at low (c) and high (d) magnification. ACE2-positive cells were observed in the brush boarder of proximal tubular cells and cytoplasm of both proximal and distal tubular cells. TMPRSS2-positive cells were observed in the cytoplasm of proximal tubular cells as well as distal tubular cells. In the glomeruli, neither ACE2 nor TMPRSS2 was expressed.

In the digestive tract, ACE2 was strongly expressed in the superficial layer of the epithelium of hypopharynx and esophagus, whereas the cytoplasm of squamous epithelium was only weakly positive for TMPRSS2 (Fig. 2A, B, 3C). In the kidney, ACE2 staining was strong in the brush boarder of proximal tubular cells, weak in the cytoplasm of proximal and distal tubular cells, and negative in the glomeruli. Expression of TMPRSS2 was strong in the cytoplasm of proximal tubular cells, weak in the distal tubular cells, and negative in the glomeruli (Fig. 4C, D).

## Discussion

In this study, we reported the immunolocalization of ACE2 and TMPRSS2, which are considered to play a pivotal role in the manifestation of COVID-19. Co-expression of ACE2 and TMPRSS2 may induce and enhance the invasion of SARS-CoV-2 into the organs ^15^. Notably, strong co-expression of ACE2 and TMPRSS2 proteins was observed in the taste buds of the tongue, nasal respiratory epithelium, trachea, bronchioles, alveoli, and the kidney. Remarkably, TMPRSS2 expression was much stronger in the peripheral alveoli than in the central alveoli. In the upper and lower respiratory tracts, the whole epithelium showed expression of ACE2 to a certain extent, whereas TMPRSS2 expression varied across sites. These results coincide with the reported clinical symptoms of COVID-19, such as loss of taste, loss of olfaction, respiratory dysfunction, and acute nephropathy ^16,17^.

The strong co-expression of ACE2 and TMPRSS2 in the taste buds may explain the high incidence of ageusia or dysgeusia in patients with SARS-CoV-2 infection. Although gene expression of ACE2 in the tongue had been examined previously ^12^, the present study is the first to reveal co-expression of ACE2 and TMPRSS2 proteins in the taste buds.

Olfaction is also impaired by SARS-CoV-2 ^6^. The current study revealed the presence of both ACE2 and TMPRSS2 in the olfactory neuroepithelium and respiratory epithelium, thus providing the first step in understanding the pathogenesis of olfactory impairment due to SARS-CoV-2 infection. Future studies to elucidate the mechanisms of sensorineural olfactory loss in COVID-19 might provide novel insights into SARS-CoV-2-related olfactory dysfunction.

The lung is an organ highly susceptible to SARS-CoV-2 infection. While ACE2 is moderately expressed in the bronchial epithelium and in type 2 pneumocytes, TMPRSS2 is strongly expressed in the cytoplasm of bronchioles and alveolar epithelial cells ^18^. Although ACE2 was found to exist on alveolar epithelial cells at approximately similar level as in the whole lung, the expression level of TMPRSS2 protein was considerably different between the peripheral and central parts of the lung. Considering the peripheral parts of the lung to strongly express TMPRSS2, along with ACE2, SARS-CoV-2 may be considered to damage the peripheral area at the very beginning of infection. Thus, the present study could histologically resolve why chest CT reveals consolidation and ground glass opacities in the bilateral peripheral lobes in confirmed COVID-19 cases ^19^. The tracheal epithelium, as well as nasal respiratory epithelium, was found to express both ACE2 and TMPRSS2. Unlike that in the nasal respiratory epithelium, TMPRSS2 was more strongly expressed in the cytoplasm of tracheal epithelium, hence suggesting the trachea to be more prone to developing clinical symptoms after SARS-CoV-2 infection than the nasal tissue.

The palate, pharynx, and esophagus are similarly lined by squamous epithelium. However, ACE2 and TMPRSS2 expression was found to be quite different across the organs in the present study. The palate displayed weak cytoplasmic staining of ACE2/TMPRSS2 in its epithelium, whereas the hypopharynx and esophagus had high expression of ACE2 and weak expression of TMPRSS2 in the epithelium. Weak-to-moderate co-expression of ACE2/TMPRSS2 in the squamous mucosa of the oropharynx and esophagus may explain the infrequent involvement of oral and throat symptoms in patients with COVID-19, without advancing to severity. Regarding ACE2 expression in the vocal cords, previous studies had provided conflicting results; while one study asserted lack of ACE2 expression in the vocal cords ^18^, the other demonstrated ACE2 expression on the surface epithelium ^20^; the present study supported the latter.

As for limitations of the present study, the evaluation was performed on young rats which may not correctly reflect the backgrounds of COVID-19 in human. Future clinical case studies and autopsy studies might strengthen this study.

## Conclusion

This study demonstrated the co-expression of ACE2 and TMPRSS2 by cells in the tongue and nasal mucosa, as well as by those in the epithelium of larynx, trachea, and bronchus. A wide range of tissues and organs are speculated to be affected by SARS-CoV-2, depending on the expression levels of ACE2 and TMPRSS2. Differential distribution of TMPRSS2 in the lung indicated COVID-19 symptoms to possibly be exacerbated by TMPRSS2 expression.

## Acknowledgement

This work was supported by JSPS KAKENHI Grant-in-Aid for Scientific Research [grant number 16K20231 and 16KT0190] and by the Smoking Research Foundation (Tokyo, Japan).

